# Network effects of the neuropsychiatric 15q13.3 microdeletion on the transcriptome and epigenome in human induced neurons

**DOI:** 10.1101/772541

**Authors:** Siming Zhang, Xianglong Zhang, Shining Ma, Carolin Purmann, Kasey Davis, Wing Hung Wong, Jonathan Bernstein, Joachim Hallmayer, Alexander E Urban

## Abstract

Heterozygous deletions in the 15q13.3 region are associated with several neuropsychiatric disorders including autism, schizophrenia, and attention deficit hyperactivity disorder. Several genes within the 15q13.3 deletion region may play a role in neuronal dysfunction, based on association studies in humans and functional studies in mice, but the intermediate molecular mechanisms remain unknown. We analyzed the genome-wide effects of the 15q13.3 microdeletion on the transcriptome and epigenome. Induced pluripotent stem cell (iPSC) lines from three patients with the typical heterozygous 15q13.3 microdeletion and three sex-matched controls were generated and converted into induced neurons (iNs) using the neurogenin-2 induction method. We analyzed genome-wide gene expression using RNA-Seq, genome-wide DNA methylation using SeqCap-Epi, and genome-wide chromatin accessibility using ATAC-Seq, in both iPSCs and iNs. In both cell types, gene copy number change within the 15q13.3 microdeletion was accompanied by significantly decreased gene expression and no compensatory changes in DNA methylation or chromatin accessibility, supporting the model that haploinsufficiency of genes within the deleted region drives the disorder. Further, we observed global effects of the deletion on the transcriptome and epigenome, with the effects being cell type specific and occurring at discrete loci. Several genes and pathways associated with neuropsychiatric disorders and neuronal development were significantly altered, including Wnt signaling, ribosome biogenesis, DNA binding, and clustered protocadherins. This molecular systems analysis of a large neuropsychiatric microdeletion can also be applied to other brain relevant chromosomal aberrations to further our etiological understanding of neuropsychiatric disorders.

## Introduction

The neuropsychiatric 15q13.3 microdeletion (OMIM 612001) is a heterozygous deletion of approximately 1.5-2 Mb at coordinates 30.5 Mb to 32.5 Mb on chromosome 15 (hg19) [1]. The deletion breakpoints lie within two regions of segmental duplications, which likely mediate the creation of the recurrent microdeletion via non-allelic homologous recombination [2]. The 15q13.3 microdeletion has an estimated prevalence of 1 in 40,000 individuals, and was first reported in 2008, in nine individuals with mental retardation, seizures, and mild facial dysmorphism [3]. Since then, several other phenotypes have been associated with this copy number variant (CNV), including autism spectrum disorder, schizophrenia, mood disorders, attention deficit hyperactivity disorder, hypotonia, and cardiac defects [1,4,5].

The most common form of the 15q13.3 microdeletion, which is the subject of our study, results in the heterozygous loss of seven protein-coding genes (CHRNA7, FAN1, TRPM1, KLF13, OTUD7A, MTMR10, ARHGAP11B) and one microRNA (MIR211) [4]. Several of these genes have previously been investigated as candidates for having a role in the neuropsychiatric symptoms associated with the CNV. The most frequently proposed candidate gene is CHRNA7, which encodes a nicotinic acetylcholine receptor and whose copy number change has been tied to reduced calcium flux in neural progenitor cells [6]. CHRNA7 expression changes in postmortem brain tissues have also been associated with schizophrenia [7], autism [8], and bipolar disorder [9]. Another possible candidate is OTUD7A, a deubiquitinase gene that in mouse models has been associated with reduced neuronal spine density, abnormal electroencephalography activity, and changes in ultrasonic vocalization and acoustic startle response [10,11]. Two other candidates genes in the deletion region are FAN1, a DNA repair nuclease that has nonsynonymous single nucleotide variants associated with both schizophrenia and autism [12,13]; and ARHGAP11B, a hominin-specific gene that increased basal progenitor cell generation in a mouse model, and that has been hypothesized to be a contributor to the evolutionary expansion of the human neocortex [14].

While previous studies have explored the role that individual genes within the 15q13.3 microdeletion may play in neuronal dysfunction, little is known about which downstream genes and molecular mechanisms are affected and how they contribute to the associated neuropsychiatric symptoms. To date, only one study has characterized the transcriptome-wide effects of the 15q13.3 microdeletion in human tissue, and this analysis was performed in non-neuronal (lymphoblastoid) cells [15]. Current knowledge is even more limited concerning the epigenetic changes associated with this CNV. Investigating whether the 15q13.3 microdeletion has genome-wide effects on different levels of genome regulation, such as DNA methylation and chromatin accessibility, will likely provide a better mechanistic understanding of this microdeletion, given that epigenetic processes play a key role in brain development and are dysregulated in a number of neuropsychiatric disorders, including major depressive disorder, autism, Rett syndrome, and schizophrenia [16].

In the present study, we analyzed the changes in expression, DNA methylation and chromatin accessibility in induced pluripotent stem cells (iPSCs) and iPSC-derived induced neurons (iNs) with the most common form of the 15q13.3 microdeletion. Due to the difficulty of accessing primary human brain tissues, iPSCs provide a useful model for studying the molecular effects of neuropsychiatric disorders. iPSCs can be reprogrammed from patient tissues and then differentiated into relevant cell types such as neurons. The use of iPSCs also allows us to follow the molecular effects of the 15q13.3 microdeletion during early neuronal development, which is not feasible with primary brain tissues. We observed genome-wide changes on all three levels of functional genomics analysis; these changes were cell type specific and many of them occurred at discrete loci. Our results revealed effects on neuropsychiatric candidate genes, gene families, and pathways linked to neuronal dysfunction, including Wnt signaling, cell adhesion, DNA binding, and protein synthesis.

## Materials and Methods

### Human Subjects

Fibroblasts derived from skin biopsies were obtained from three patients with the most common 15q13.3 microdeletion and three sex-matched controls (Fig. 1a). Samples were obtained after informed consent was given by all subjects and were subsequently de-identified. All procedures were approved by the Stanford University Institutional Review Board.

**Figure 1.**
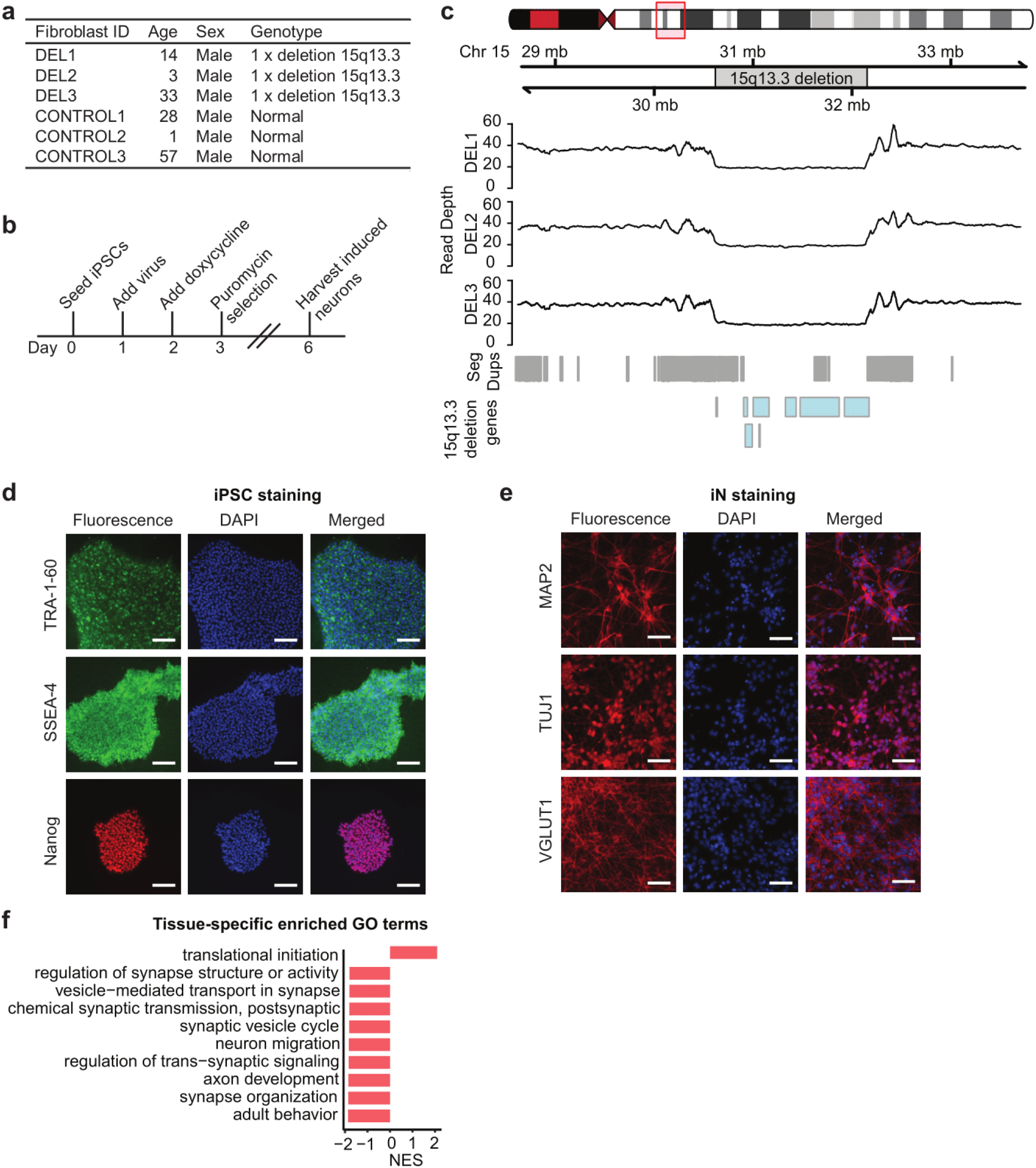
Generation and characterization of CNV lines. (a) Information on donors of fibroblasts, including age, sex, and genotype. (b) Induced neuron generation timeline. (c) Sequencing read depth of fibroblast samples. (d) Immunocytochemistry for the pluripotency markers Nanog, TRA-1-60, and SSEA-4 on iPSCs (scale bar: 200 μm). (e) Immunocytochemistry for the neuronal markers MAP2, TUJ1 and VGLUT1 on day 6 induced neurons (scale bar: 200 μm). (f) Significant gene ontology biological process terms from gene set enrichment analysis of genes differentially expressed between iPSCs and iNs (top 10 based on normalized enrichment score)

### Generation of iPSCs and induced neurons (iNs)

iPSCs were generated from fibroblasts using Sendai viral vectors from the CytoTune-iPS 2.0 Sendai Reprogramming Kit (Life Technologies, Carlsbad, CA, USA). Induced neurons (iNs) were differentiated from iPSCs using the neurogenin-2 induction method [17]. Day 6 iNs were collected for downstream analysis (Fig. 1b). See Supplementary Information for further details.

### Immunocytochemistry

iPSCs were stained with the pluripotency markers Nanog, Tra-1-60, and SSEA-4. Day 6 iNs were stained with the neuronal identity markers TUJ1, VGLUT1, and MAP2. See Supplementary Information for further details.

### DNA sequencing

Genomic DNA was isolated from fibroblasts using the Quick-DNA Universal Kit (Zymo Research, Irvine, CA, USA). DNA libraries were generated using the TruSeq DNA Nano Library kit (Illumina, San Diego, CA, USA) and sequenced on the Illumina HiSeq X using 2×150 bp paired-end runs. Sequencing reads were mapped to GRCh37 using BWA [18]. BEDtools [19] was used to convert genomic coordinates to GRCh38 and calculate read depth coverage within the 15q13.3 microdeletion region.

### RNA sequencing

RNA libraries were prepared using the NEBNext Ultra Directional RNA Library preparation kit (New England Biolabs, Ipswich, MA, USA) and sequenced on the Illumina NextSeq 500. Reads were mapped to GRCh38 using Tophat [20] and differential expression analysis was performed using DESeq2 [21]. WebGestalt and Ingenuity Pathway Analysis (QIAGEN Inc.) were used for gene set enrichment analysis and over-representation analysis [22]. See Supplementary Information for further details.

### Genome-wide targeted-capture DNA methylation sequencing

Libraries for DNA methylation were prepared using the SeqCap Epi System (Roche, Basel, Switzerland) and sequenced on the Illumina HiSeq 4000. Reads were aligned to GRCh38 using Bowtie 2 [23], DNA methylation ratios for CpGs were called by Bismark [24], and differentially methylated regions (DMRs) were identified using Metilene [25]. Gene set enrichment analysis was performed on lists of DMR-associated genes using WebGestalt [22]. See Supplementary Information for further details.

### Assay for transposase-accessible chromatin (ATAC) sequencing

ATAC-Seq libraries were generated as previously described [26] and sequenced on the Illumina NextSeq 500. Reads were mapped to GRCh38 with Bowtie2 [23] and peaks were called using MACS2 [27]. Differential chromatin accessibility analysis was performed using Diffbind [28]. Homer motif enrichment analysis [29] was performed using a list of differentially open promoters and a custom motif file, generated as previously described [30]. See Supplementary Information for further details.

### Correlation analysis

We calculated the pairwise correlations between gene expression log2 fold change, mean methylation difference, and chromatin accessibility log2 fold change, using genes that were significant (padj <= 0.05) in at least one of the data sets within each pairwise comparison. For the Methyl-Seq and ATAC-Seq analysis, only genes with significant DMRs or peaks in the promoter (region from transcriptional start site to 2 kb upstream) were included.

## Results

### Characterization of 15q13.3 iPSCs and iNs

To study the molecular effects of the 15q13.3 microdeletion, we carried out genome-wide analyses of patterns of gene expression, DNA methylation, and chromatin accessibility in iPSCs and iNs from three patients with the 15q13.3 microdeletion and three sex-matched controls. The presence of heterozygous deletions at the 15q13.3 locus in the patient fibroblast samples was confirmed using read depth coverage from whole genome sequencing (Fig. 1c). The deletions in all three samples had similar breakpoints and were all approximately 1.5 Mb in size, the most common deletion size found in 15q13.3 deletion patients. iPSCs were generated from fibroblast lines using the CytoTune-iPS Sendai Reprogramming Kit and stained positive for the pluripotency markers SSEA4, TRA-1-60, and Nanog (Fig. 1d). Induced neurons (iNs) were generated from iPSCs via the neurogenin-2 induction method. Day 6 iNs exhibited neurite morphology and stained positive for the neuronal markers TUJ1, VGLUT1, and MAP2 (Fig. 1e). Gene set enrichment analysis also showed that genes involved in neuronal processes were up-regulated in iNs compared with iPSCs (Fig. 1f).

### Transcriptional impact of the 15q13.3 microdeletion

We performed RNA-Seq on six 15q13.3 deletion clonal lines (3 individuals, 2 clones per individual) and six control clonal lines (3 individuals, 2 clones per individual) at the iPSC and iN stages to determine the effects of the heterozygous 15q13.3 microdeletion on expression both within the deletion and genome-wide. We first looked for evidence of haploinsuffiency or dosage compensation within the 15q13.3 microdeletion, which contains seven protein-coding genes. In iPSCs, five genes, KLF13, MTMR10, OTUD7A, FAN1, and ARHGAP11B, showed significantly decreased gene expression in 15q13.3 deletion cases (padj <= 0.05). Additionally, CHRNA7 was expressed at very low levels and was non-significantly decreased in deletion cases (Fig. 2a). In iNs, six genes, KLF13, MTMR10, OTUD7A, FAN1, ARHGAP11B, and CHRNA7, were significantly decreased in 15q13.3 deletion cases (padj <= 0.05). The gene TRPM1 was not detected above threshold levels in iPSCs or iNs (Figs. 2a, b). We also analyzed the gene expression data set for a position effect, in which genes located near but outside the 15q13.3 microdeletion are affected. In both iPSCs and iNs, no significant enrichment was observed for genes within 5 Mb from the 15q13.3 deletion. Using Webgestalt, we carried out an analysis to determine if any cytogenetic bands were enriched in differentially expressed genes. As expected, the 15q13.3 locus was enriched in both iPSCs and iNs (padj <= 0.05). In addition, the locus 10q11.2 was significantly enriched in iPSCs. One of the differentially expressed genes within this region is GPRIN2, which regulates neurite outgrowth and has been identified as being affected by copy number variation in several patients with autism [31].

**Figure 2.**
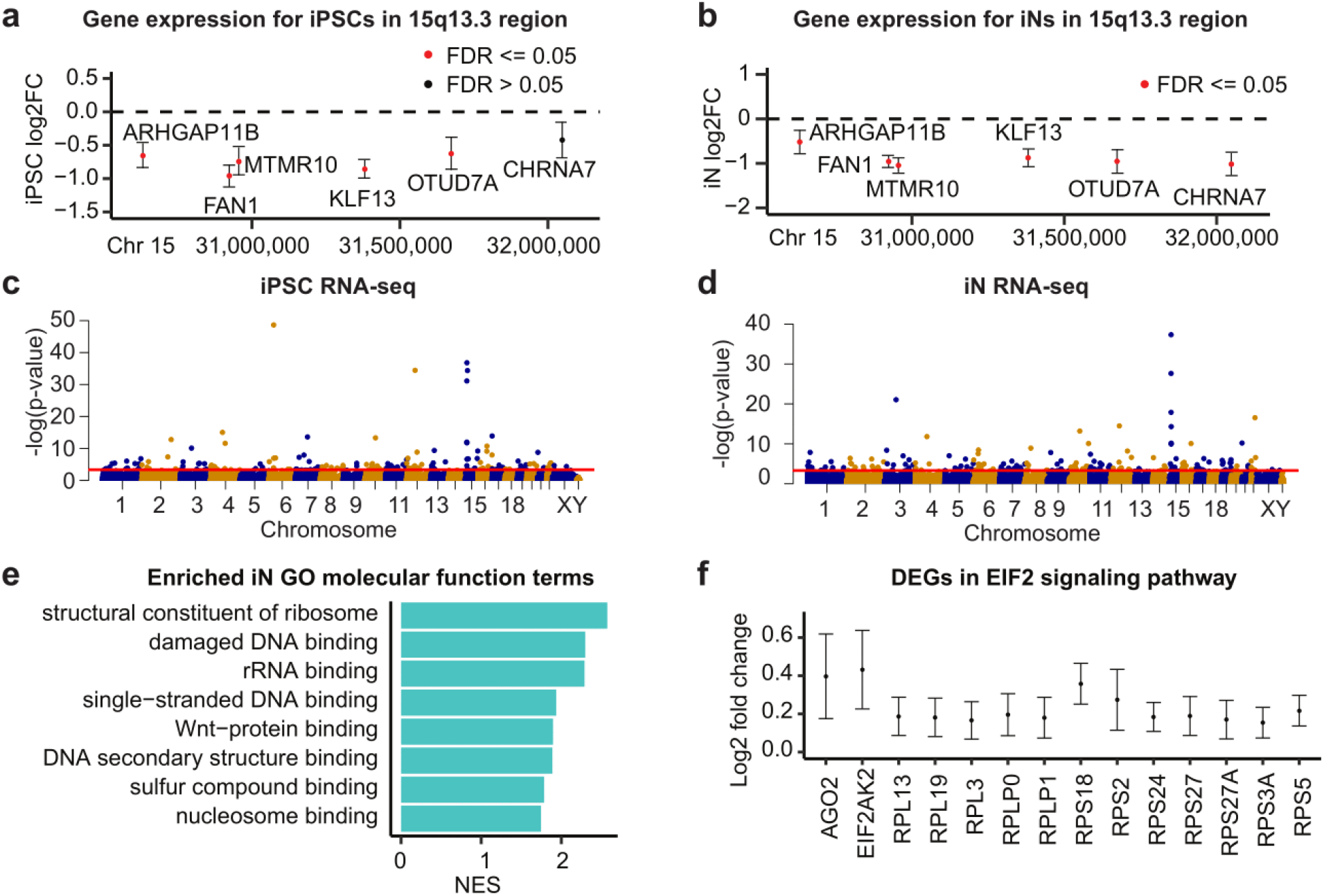
Gene expression changes within the 15q13.3 microdeletion and genome-wide. (a, b) Log2 fold change of protein-coding 15q13.3 genes in deletion samples compared with controls in iPSCs and iNs (95% confidence interval shown). (c, d) Manhattan plots of RNA-Seq genes in iPSCs and iNs, with the red threshold line indicating 0.05 FDR significance. (e) Significant gene ontology molecular function terms from gene set enrichment analysis of differentially expressed iN genes (padj <= 0.05), sorted by normalized enrichment score. (f) Log2 fold change of EIF2 signaling genes differentially expressed in deletion iNs compared with control iNs (95% confidence interval shown).

We observed genome-wide changes in the expression of genes as a result of the 15q13.3 microdeletion. 178 genes were differentially expressed (DE) between 15q13.3 deletion cases and controls at the iPSC stage and 369 genes were DE at the iN stage (Figs. 2c, d and Supplementary Tables S1, S2). The much larger number of DE genes at the iN level compared with the iPSC level suggests the 15q13.3 microdeletion may have stronger downstream effects in a more relevant cell type with regard to the associated neuropsychiatric phenotypes. Out of 41 genes overlapping between the iPSC and iN DE gene lists, all showed the same direction of fold change in gene expression except for S100A10, which shifted from decreased expression in deletion cases for iPSCs to increased expression in deletion cases for iNs. S100A100, a member of the S-100 family of calcium binding proteins, is involved in depression-like behaviors through regulation of serotonin receptor activity [32]. At the iN stage, a number of genes were identified as differentially expressed that have been previously implicated in neuropsychiatric disorders associated with the 15q13.3 microdeletion. These genes include VIPR2, PRODH, and DGCR6 for schizophrenia [33–35]; CACNG3, SCN8A, SPATA5, and KCNA2 for epilepsy [36–39]; and LINS1 and DCPS for intellectual disability [40–42].

Using the lists of differentially expressed genes (padj <= 0.05), we performed gene set enrichment analysis using Webgestalt. In iPSCs, no gene ontology molecular function terms were significantly enriched. In iNs, we identified several significant gene ontology molecular function terms, with the top terms grouping into the categories of ribosome function, nucleic acid binding, Wnt binding, and sulfur compound binding (Fig. 2e). In addition, we performed over-representation analysis using Ingenuity Pathway Analysis (IPA) to identify enriched canonical pathways. Within iNs, there was an enrichment of the EIF2 signaling pathway, which is involved in translational control (padj <= 0.05). The genes driving this enrichment, which were all significantly up-regulated in 15q13.3 deletion samples, included AGO2, which plays a key role in RNA interference; EIF2AK2, a kinase that phosphorylates translation initiation factor EIF2S1; and 12 ribosomal protein genes (Fig. 2f).

### DNA methylation analysis

We performed genome-wide targeted-capture bisulfite sequencing on the same cohort of cell lines used for gene expression analysis (six 15q13.3 deletion clonal lines and six control clonal lines at the iPSC and iN stages). One iN control line was excluded from final analysis due to clustering with iPSCs rather than iNs. Within the 15q13.3 microdeletion, no differentially methylated regions (DMRs) were identified in iPSCs or iNs with the exception of one DMR located in the gene body of non-coding RNA LINC02352. The absence of significant DNA methylation changes in the 15q13.3 genes was consistent with the lack of dosage compensation seen in the gene expression data. In addition, we did not observe an enrichment in DMRs within 5 Mb of the 15q13.3 microdeletion.

Genome-wide, 416 DMRs were identified in iPSCs and were annotated to 147 promoter regions, 404 gene bodies, and 77 intergenic regions (Figs. 3a, c and Supplementary Table S3). At the iN level, 408 DMRs were identified and annotated to 160 promoter regions, 277 gene bodies, and 80 intergenic regions (Figs. 3b, d and Supplementary Table S4). After assigning each gene to a unique DMR, we obtained a list of 355 differentially methylated genes in iPSCs and 365 differentially methylated genes in iNs. The effect direction for the mean methylation difference between cases and controls was concordant for 215 out of 216 genes differentially methylated in both iPSCs and iNs, suggesting a high degree of similarity in DNA methylation patterns between the two tissues. The one exception that was non-concordant was a gene associated with two different DMRs in the two tissue types.

**Figure 3.**
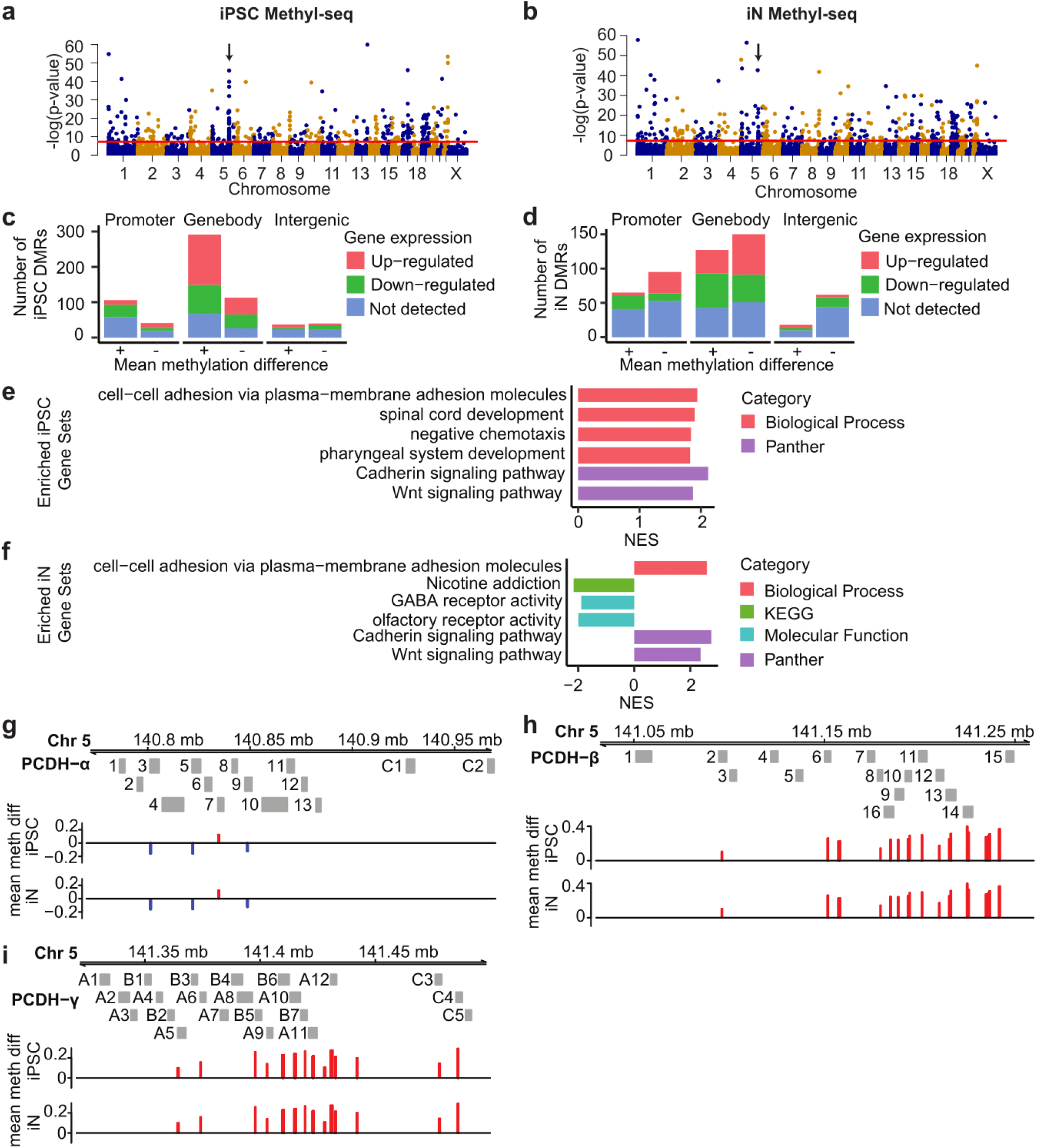
Differentially methylated regions (DMRs) in 15q13.3 deletion versus control cell lines. (a, b) Genome-wide distribution of DMRs in iPSCs and iNs. DMRs above red threshold line have >=0.1 mean methylation difference and padj <= 0.05. Arrows indicate PCDH gene clusters. (c, d) DMRs in iPSCs and iNs, split by DMR location, direction of mean methylation difference and direction of expression change in nearest gene. (e, f) Significant terms from gene set enrichment analysis of differentially methylated genes in iPSCs and iNs, sorted by normalized enrichment score. (g, h, i) DMRs located near the (g) alpha, (h) beta and (i) gamma PCDH families. DMRs are colored red for higher mean methylation in deletion samples and blue for lower.

We performed gene set enrichment analysis using the lists of DMR-associated genes. In both iPSCs and iNs, we observed an enrichment in the gene ontology biological process (GOBP) category of cell-cell adhesion via plasma-membrane adhesion molecules (GO:0098742). Under the category of PANTHER pathways, both the cadherin signaling pathway (P00012) and the Wnt signaling pathway (P00057) were enriched in iPSCs and iNs (Figs. 3e, f). The enriched PANTHER and GOBP terms were mainly driven by a large number of DMRs located near protocadherins (PCDHs) (Figs. 3g, h, i), which are cell-adhesion proteins primarily expressed in the developing nervous system [43]. PCDHs have been directly implicated in several neuronal disorders, including epilepsy, autism, and schizophrenia [44]. Specifically, differential methylation of PCDHs has been observed in a number of neurodevelopmental disorders, including Down syndrome, Williams-Beuren syndrome, 7q11.23 duplication syndrome, and depression [45]. Besides the protocadherin-related pathways, we also observed an enrichment in two other neuronally related categories within iNs: the KEGG category of nicotine addition (hsa05033) and the gene ontology molecular function category of GABA receptor activity (GO:0016917) (Fig. 3f).

### Chromatin accessibility analysis

We examined chromatin accessibility in three 15q13.3 deletion lines (3 individuals, 1 clone per individual) and three control lines (3 individuals, 1 clone per individual) at the iPSC and iN stage. In iPSCs, the average chromatin accessibility log2 fold change was −1.039 within the 15q13.3 region and 0.011 genome-wide. In iNs, the average chromatin accessibility log2 fold change was −0.905 within the 15q13.3 region and −0.003 genome-wide. In both cases, this represented the loss of one copy number within the 15q13.3 deletion region. Within 5 Mb of the deletion, we did not observe any enrichment in differentially accessible peaks.

Genome-wide, we identified 50 differentially accessible peaks in iPSCs and 72 peaks in iNs (Figs. 4a, b and Supplementary Tables S5, S6). These peaks were annotated to 9 promoter regions, 21 gene bodies, and 22 intergenic regions in iPSCs and to 14 promoter regions, 23 gene bodies, and 40 intergenic regions in iNs (Figs. 4c, d). After assigning each gene to a unique peak, we identified 45 differentially accessible genes in iPSCs and 68 differentially accessible genes in iNs. Five genes were differentially accessible in both tissues, all with a concordant direction of effect for the chromatin accessibility change between cases and controls.

**Figure 4.**
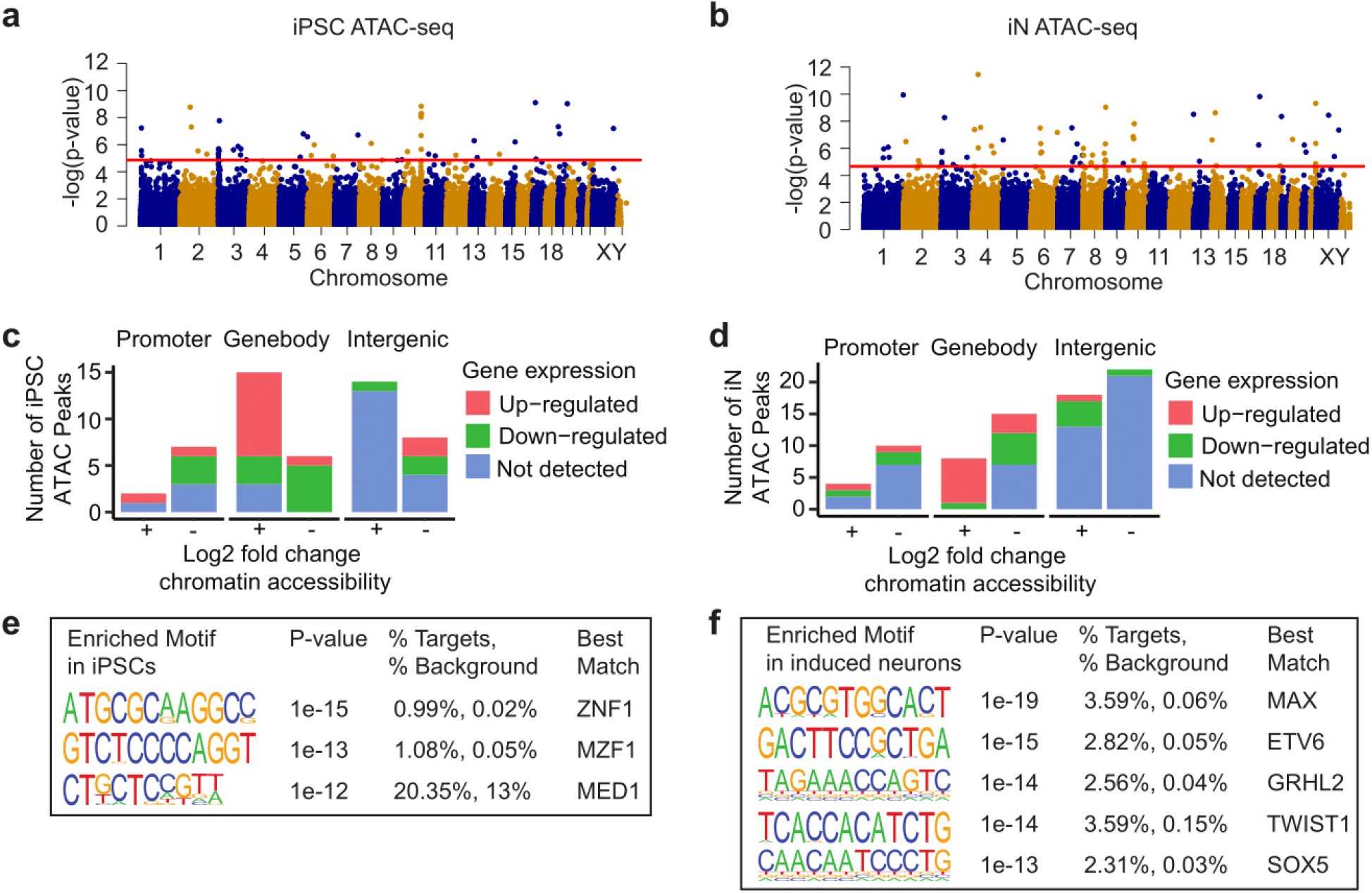
Differential chromatin accessibility peaks in 15q13.3 deletion versus control cell lines. (a, b) Genome-wide distribution of peaks identified from ATAC-Seq in iPSCs and iNs. Peaks above red threshold line have padj <=0.05. (c, d). Differentially accessible ATAC-Seq peaks in iPSCs and iNs, split by peak location, direction of chromatin accessibility change and direction of expression change in nearest gene. (e, f). Enriched motifs in iPSCs and iNs from de novo Homer analysis with p-value <= 10^−12^, best match transcription factor expressed in the same tissue, and best match score >= 0.6.

We next examined transcription factor binding sites (TFBSs) enriched in a list of differentially open promoters, generated as previously described [30]. In iPSCs, three enriched motifs (pval<=10-12) were associated with transcription factors that were expressed in the same tissue: ZNF711, MZF1, and MED1 (Fig. 4e). In iNs, five expressed transcription factors associated with enriched motifs (pval<=10-12) were similarly identified: MAX, ETV6, GRHL2, TWIST1, and SOX5 (Fig. 4f). Out of the eight transcription factors associated with enriched motifs across iPSCs and iNs, four (ZNF711, MZF1, MAX, SOX5) have been previously implicated in neuropsychiatric disorders. Variants in ZNF711 and MZF1 have been associated with non-syndromic intellectual disability and Alzheimer’s disease, respectively [46,47]. Genetic variants in SOX5 have been associated with a range of neuropsychiatric disorders, including major depressive disorder [48], amyotrophic lateral sclerosis [49], Alzheimer’s disease [50] and intellectual disability [51]. Lastly, the transcription factor MAX has been linked to depression-like behaviors in mouse models [52] and has been shown to be down-regulated in the dorsolateral prefrontal cortex of schizophrenia patients [53].

### Correlation and overlap analysis

To determine if the RNA-Seq, Methyl-Seq and ATAC-Seq omics data sets exhibited a linear relationship, we calculated pairwise Pearson correlations between transcription, promoter-associated DNA methylation and promoter-associated chromatin accessibility. A weak positive correlation was seen between transcription and chromatin accessibility in both iPSCs (R=0.21, p=0.019) and iNs (R=0.27, p=2.5×10^−6^) (Figs. 5a, d). A weak negative correlation was seen between transcription and methylation in iNs (R=− 0.22, p =1.4×10^−5^) (Fig. 5e). The correlation between transcription and methylation in iPSCs was also negative but was nonsignificant (R=−0.14, p=0.053) (Fig. 5b). A strong negative correlation was observed between methylation and chromatin accessibility in iPSCs (R=−0.67, p<2.2×10^−16^) and iNs (R=−0.64, p<2.2×10^−16^) (Figs. 5c, f). These trends were consistent with the expected positive correlation between chromatin accessibility and transcription, the expected negative correlation between DNA methylation and transcription, and the expected negative correlation between DNA methylation and chromatin accessibility.

**Figure 5.**
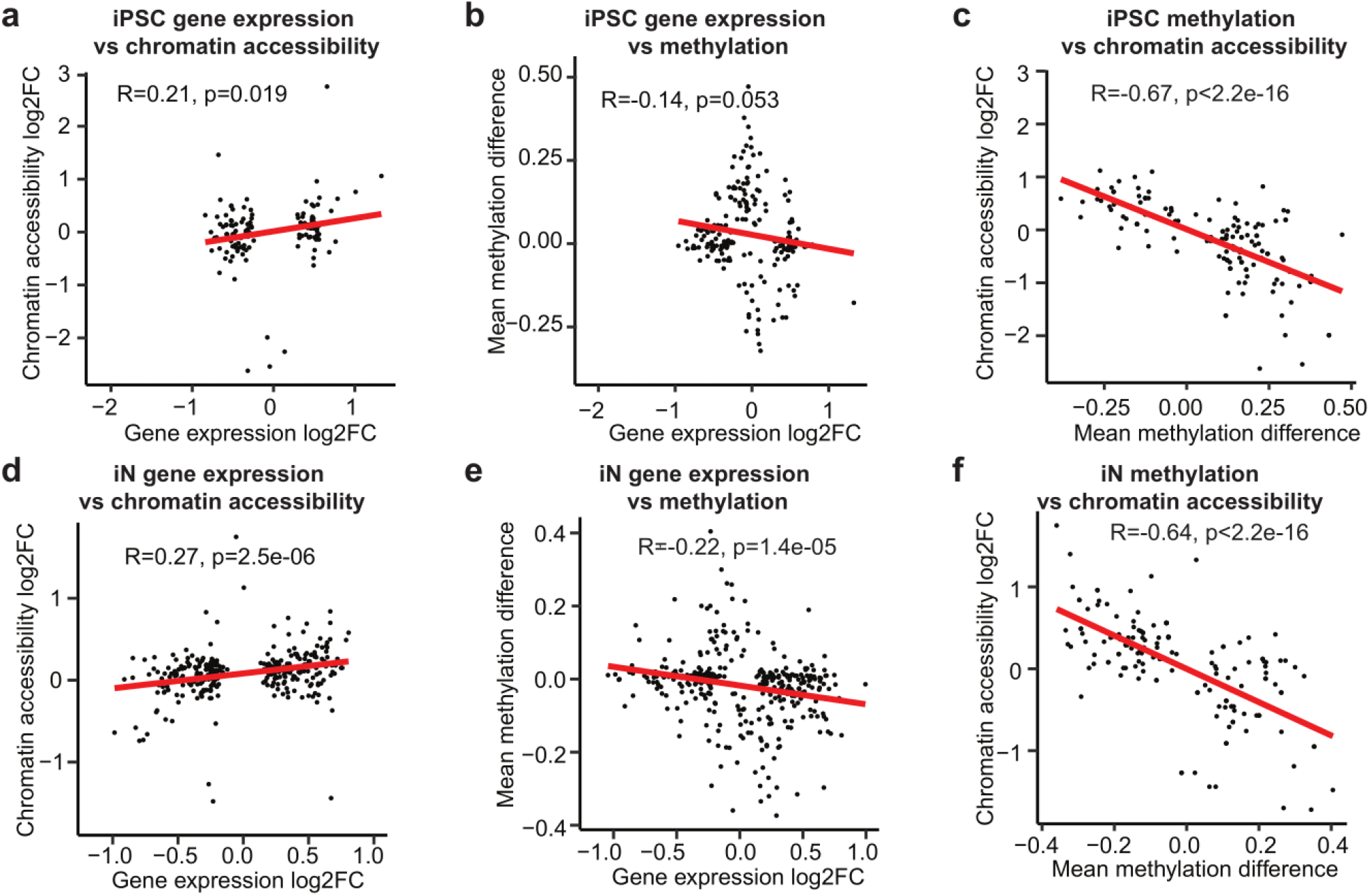
Multi-omics correlation analysis. Correlation between (a, b) gene expression log2 fold change and chromatin accessibility log2 fold change, (c, d) gene expression log2 fold change and mean methylation difference and (e, f) mean methylation difference and chromatin accessibility log2 fold change.

We also calculated pairwise correlations between transcription, DNA methylation and chromatin accessibility, between the two tissue types. Between transcription and chromatin accessibility, no significant correlation was observed (Supplementary Figs. S1d, h). Between transcription and DNA methylation, a very weak negative correlation was observed (Supplementary Figs. S1e, i). A moderate to strong negative correlation was seen between methylation and chromatin accessibility between different tissue types (Supplementary Figs. S1f, g). Finally, we examined the correlation between tissue types within each omics analysis, e.g. between iPSC transcription and iN transcription. We observed a moderate positive correlation for transcription (R=0.56, p<2.6×10^−16^), a very strong positive correlation for DNA methylation (R=0.92, p< 2.2×10^−16^), and a very strong positive correlation for chromatin accessibility (R=0.84, p< 2.6×10^−5^) (Supplementary Figs. S1a, b, c).

Despite the significant linear correlation between the three omics data sets within each tissue type, the overlap in significant genes was very limited. Only 15 genes were significant across at least two omics data sets in iPSCs and 13 genes were significant across at least two data sets in iNs. A single gene, KCNH8, was significant in all three omics analyses for iNs. KCNH8 encodes a voltage gated potassium channel that is primarily expressed in the nervous system [54]. We also identified four genes that were significant in multiple omics data sets in both iPSCs and iNs. Interestingly, all four have been implicated in studies of neuropsychiatric disorders. These genes were CNTN4, a cell adhesion molecule associated with autism [55] and schizophrenia [56]; PCDHGA10, a member of the protocadherin family of cell adhesion proteins which has been implicated in schizophrenia and autism [44]; SLC39A4, a zinc transport protein known to be regulated by SHANK3, a highly penetrant risk factor for autism [57]; and SORCS1, which alters amyloid precursor protein processing and is associated with Alzheimer’s disease [58].

## Discussion

Our multi-omics analysis of iPSCs and iPSC-derived iNs carrying the 15q13.3 microdeletion allowed us to interrogate the global molecular impact of a large CNV that is strongly associated with a range of neuropsychiatric disorders. Within the 15q13.3 microdeletion, the genes CHRNA7, KLF13, MTMR10, OTUD7A, FAN1, and ARHGAP11B were expressed in both iPSCs and iNs. The remaining protein-coding gene in the region, TRPM1, was not expressed above threshold levels in iPSCs or iNs, which is in agreement with previous expression data suggesting it is primarily expressed in the retina [59]. In both iPSCs and iNs, all expressed genes had decreased expression in deletion samples, suggesting simple dosage sensitivity as a mechanism of action. We did not see significant changes in DNA methylation or chromatin accessibility within the deletion after accounting for the change in copy number, which was consistent with the lack of dosage compensation observed in the gene expression data. We also did not observe a position effect, in which genes near (within 5 Mb) but outside the CNV are affected.

We saw genome-wide changes in the expression, DNA methylation and chromatin accessibility patterns of iPSCs and iPSC-derived neurons as a result of the 15q13.3 microdeletion. In the pairwise correlation analyses of the omics data sets, DNA methylation had a negative correlation with both transcription and chromatin accessibility, whereas transcription and chromatin accessibility were positively correlated. The Pearson correlation between DNA methylation and chromatin accessibility was much stronger than their respective correlations with transcription. This trend held true even when comparing DNA methylation and chromatin accessibility across different tissues, and was likely due to the greater stability of the epigenome compared with gene expression. Both the direction and relative strength of correlation between these three omics layers were consistent with a recent study that profiled transcription, methylation and accessibility in mouse embryonic stem cells. In the promoter-associated pseudo-bulk data from that study, DNA methylation-accessibility had the highest correlation coefficient, followed by DNA methylation-transcription and accessibility-transcription [60].

At the gene expression level, our analysis in iNs compared with the analysis in iPSCs revealed a greater number of changes in neuropsychiatric-related genes and pathways. Among the list of differentially expressed iN genes, two of particular interest were PRODH and DGCR6, which are major candidate genes in the 22q11.2 microdeletion, another large CNV like the 15q13.3 microdeletion that is strongly associated with increased risk of neuropsychiatric disorders [61]. From the gene set enrichment analysis, the gene ontology molecular function categories enriched in iNs included a number of processes tied to abnormal neurodevelopment, such as Wnt binding, ribosome biogenesis, and DNA binding. Wnt signaling plays an important role in a number of neurodevelopmental and neurophysiological processes, such as synapse assembly, neuronal generation and differentiation, and glutamatergic and GABAergic neurotransmission [62]. Changes in the expression of Wnt pathway components are associated with several neuronal disorders including schizophrenia, bipolar disorder, Alzheimer’s disease, and autism [63]. Similarly, the dysregulation of ribosome biogenesis has been implicated in neuropsychiatric disorders in previous studies of postmortem brains [64,65], olfactory neurosphere derived cells [66], and neural progenitor cells [67]. Another enriched gene ontology category, damaged DNA binding, is also known to play a key role in neuropsychiatric disorders. Mutations in DNA repair genes have been linked to schizophrenia and autism [68], and oxidative stress, a major trigger of DNA damage, has been associated with a number of neuropsychiatric disorders including major depressive disorder, bipolar disorder, and schizophrenia [69].

In our analysis of the epigenome, we also identified an enrichment in genes connected to Wnt signaling, although the specific genes involved did not significantly overlap with those affected at the gene expression level. In the DNA methylation analysis, the top PANTHER pathways enriched in both iPSCs and iNs were cadherin signaling and Wnt signaling. The genes differentially methylated in these pathways were primarily composed of protocadherins, a group of cell adhesion genes known to modulate the Wnt signaling pathway [70,71]. From the chromatin accessibility analysis, we also identified several transcription factors with known connections to Wnt signaling, including MZF1, whose transcription factor binding sites are enriched in promoters of Wnt pathway genes [72]; TWIST1, which is activated by canonical Wnt signaling [73]; and SOX5, which is associated with WNT signaling activity in human SH-SY5Y neuroblastoma cells [50].

In addition to the convergence on the Wnt signaling pathway, our multi-omics analysis identified four genes (CNTN4, PCDHGA10, SLC39A4, SORCS1) enriched in multiple omics data sets in both iPSCs and iNs. All four genes have been previously associated with schizophrenia, autism, or Alzheimer’s disease. This enrichment of previously reported neuropsychiatric associated genes and pathways in the omics overlap analysis suggests that integrating multiple omics layers could help us narrow down on candidate genes and molecular networks for CNV-associated psychiatric disorders like the 15q13.3 microdeletion syndrome, which often have a high degree of variability at the phenotypic and likely molecular level.

Follow-up studies can now be conducted that will use more than the single neuronal cell type used here, which were early-stage induced neurons that can mature to resemble excitatory projection neurons. Given the diverse range of phenotypes associated with the 15q13.3 microdeletion, it would be beneficial in future studies to look at other neuronal cell types, which may be affected differently by the CNV, as well as their interactions with glial cells also carrying the microdeletion.

In conclusion, we generated iPSCs and iNs with the most common variant of the 15q13.3 microdeletion, which provide a useful tool to study this large CNV as well as acting as an entry point for understanding the molecular etiology of major neuropsychiatric disorders in general. Our analysis of these cell lines showed that all expressed genes within the CNV region are down-regulated following the change in DNA copy number, supporting the model that haploinsufficiency of genes within the deleted regions forms the molecular basis of the observed phenotypes. Our analyses of gene expression, DNA methylation and chromatin accessibility revealed downstream changes that were both global in nature and affecting discrete loci, genes, gene families and pathways, such as Wnt signaling, ribosome function, DNA binding, and cell adhesion, suggesting these molecular pathways may be disrupted in 15q13.3 microdeletion patients during brain development and brain function. Beyond the analysis of the 15q13.3 microdeletion, our study provides a blueprint for assaying the molecular consequences of large neuropsychiatric chromosomal aberrations in general. The many points of convergence with loci already associated with normal and abnormal brain function underline how the study of variants such as those on chromosome 15q13.3 can inform our general understanding of the molecular basis of neuropsychiatric disorders.

## Acknowledgements

We would like to thank Guangwen Wang from the Stanford Stem Cell Core for the generation of iPSCs and Javier Fernandez Alcudia from the Stanford Gene Vector and Virus Core for the production of lentiviruses for neurogenin-2 induction.

This work was supported by the National Science Foundation (SZ), NIH grants 5P50HG007735-05 (AEU, WHW), 1R01HG010359-01A1 (WHW), 1DP2MH100010-01 (AEU), and March of Dimes Foundation Research Grant #6-FY13-142 (AEU). AEU is a Tashia and John Morgridge Faculty Scholar at the Stanford Child Health Research Institute.

## Author Contributions

SZ and AEU designed the study. SZ, XZ, SM, and WHW developed the analysis pipeline. SZ, JB, JH, CP and KD contributed to cell line generation and characterization. SZ performed the sequencing experiments and carried out the data analyses, with contributions from SM. SZ and AEU wrote the manuscript.

## Availability of data

The data sets generated and analyzed in the current study are available at NCBI GEO GSE135131 (token whobqkaghtyjbkf) and NCBI SRA PRJNA557485.

