## Supplementary Figure 1 for "Network effects of the neuropsychiatric 15q13.3 microdeletion on the transcriptome and epigenome in human induced neurons"

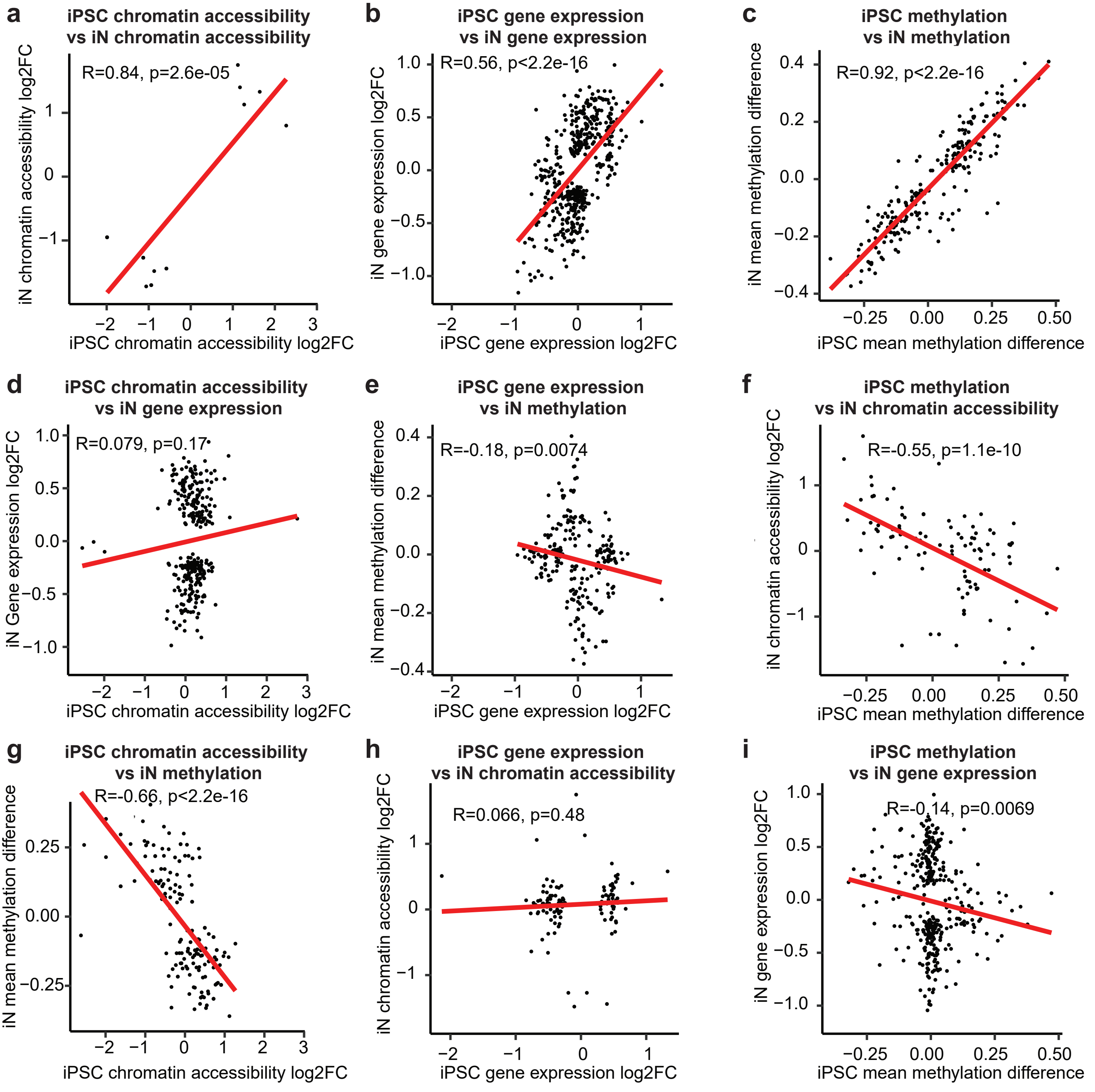


**Supplementary Figure S1. Multi-omics correlation analysis between different cell types.** Correlation between iPSCs and iNs for (a) chromatin accessibility log2 fold change, (b) gene expression log2 fold change and (c) mean methylation difference. (d-i) Correlation between iPSCs and iNs across pairwise combinations of gene expression, DNA methylation, and chromatin accessibility.
