## Supplementary Methods for "Network effects of the neuropsychiatric 15q13.3 microdeletion on the transcriptome and epigenome in human induced neurons"

**Generation of iPSCs and induced neurons**

iPSCs were generated from fibroblasts using Sendai viral vectors Oct4, Sox2, Klf4, and c-Myc from the CytoTune-iPS 2.0 Sendai Reprogramming Kit (Life Technologies, Carlsbad, CA, USA). iPSCs were maintained on irradiated mouse embryonic fibroblasts. Individual iPSC colonies were manually selected between 3-4 weeks following Sendai virus transduction for further expansion as clonal lines. The expanded iPSCs were grown on Matrigel (Corning, Corning, NY, USA) coated plates in mTeSR1 media (Stem Cell Technologies, Vancouver, Canada) and passaged using 0.02% EDTA (Lonza, Basel, Switzerland). All cell lines were confirmed to be free of mycoplasm contamination.

For each patient and control subject, two clonal iPSC lines were selected for neural differentiation using the neurogenin-2 induction method [17]. On day 0, iPSCs were dissociated into single cells using Accutase (Thermo Fisher Scientific, Waltham, MA, USA). On day 1, the lentivirus vectors FUW-rtTA, which expresses rtTA, and pTet-O-Ngn2-puro, which expresses Ngn2 and the puromycin resistant gene under control of the TetON promoter, were added to mTeSR1 medium containing 10μM rock inhibitor (Tocris Bioscience, Bristol, United Kingdom). On day 2, the culture medium was replaced with DMEM/F12 (Life Technologies) mixed with N2 supplement (Life Technologies), penicillin-streptomycin (Life Technologies), and doxycycline at 2ug/ml (Sigma Aldrich, St. Louis, MO, USA) to induce TetO expression. On days 3-5, puromycin at 2ug/ml (Sigma Aldrich) was added to the media to select for cells expressing Ngn2. On day 6, early stage induced neurons (iNs) were collected for downstream analysis (Fig. 1b).

**Immunocytochemistry**

To confirm successful reprogramming, iPSCs were stained with the pluripotency markers Nanog, Tra-1-60, and SSEA-4. Day 6 iNs were stained for the neuronal identity markers TUJ1, VGLUT1, and MAP2. For both iPSCs and iNs, cells were fixed with 4% paraformaldehyde for 15 minutes, permeabilized with 0.1% Triton X-100 for 10 minutes, blocked with 5% goat serum for one hour and labeled with primary antibody overnight at 4℃. The samples were incubated for one hour with the secondary antibody, followed by DAPI staining for 10 minutes. The following primary antibodies were used: mouse SSEA4 (ST11015, ESI Bio, Alameda, CA, USA), mouse TRA-1-60 (ST11016, ESI Bio), rabbit Nanog (09-0020, Stemgent, Cambridge, MA, USA), rabbit TUJ1 (1-15-56, Biolegend, San Diego, CA, USA), rabbit VGLUT1 (135 302, Synaptic Systems, Goettingen, Germany), and mouse MAP2 (M1406, Sigma Aldrich). The following secondary antibodies and stains were used: Alexa Fluor 488 goat anti-mouse (A11029, Life Technologies), Alexa Fluor 555 goat anti-mouse (A21422, Life Technologies), Alexa Fluor 555 goat anti-rabbit (A21428, Life Technologies), and DAPI (D9542, Sigma Aldrich). All primary antibodies were used at 1:100 dilution and all secondary antibodies were used at 1:1000 dilution. Cells were imaged using the Leica AF 6000 microscope at 20x magnification using the same acquisition settings for all samples and analyzed using ImageJ.

**RNA sequencing**

Total RNA was extracted from samples using the Direct-zol RNA MiniPrep Kit (Zymo Research, Irvine, CA, USA) and mRNA was isolated using the Dynabeads mRNA Purification Kit (Thermo Fisher Scientific). Libraries were prepared using the NEBNext Ultra Directional RNA Library preparation kit (New England Biolabs, Ipswich, MA, USA) and sequenced on the Illumina NextSeq 500 using 2x150 bp paired-end runs with 1% PhiX spike-in. The sequencing reads were mapped to GRCh38 using Tophat [20]. Principal component analysis and differential gene expression analysis were performed using DESeq2 [21], with significant genes defined as having padj <= 0.05. WebGestalt and Ingenuity Pathway Analysis (Qiagen, Hilden, Germany) were used for gene set enrichment analysis and over-representation analysis [22].

**Genome-wide targeted-capture DNA methylation sequencing**

Genomic DNA was extracted from samples using the Quick-DNA Universal Kit (Zymo Research). Bisulfite converted libraries were prepared using the SeqCap Epi CpGiant System (Roche, Basel, Switzerland), which targets 80.5 Mb containing over 5.5 million CpGs. Libraries were sequenced on the Illumina HiSeq 4000 using 2x150 bp paired-end runs and 30% PhiX spike-in. Sequencing reads were trimmed using Cutadapt and mapped to GRCh38 using Bowtie 2 [23]. DNA methylation ratios for CpGs, defined as methylated reads divided by total reads, were called by Bismark [24]. The iN sample for CONTROL1-1 showed up as an outlier in clustering analysis, grouping with the iPSCs, and was removed from subsequent analysis. Differentially methylated regions (DMRs) were identified using Metilene [25] using a minimum cutoff of 0.1 mean methylation difference between deletion and control samples and padj <= 0.05. For association with nearby genes, DMRs were assigned first to promoter regions (defined as the 2 kb region upstream of the transcriptional start site), then to gene bodies, and finally to the nearest gene (intergenic). For genes with multiple significant DMRs, the DMR with the most significant p-value was chosen for downstream gene-based analyses. Gene set enrichment analysis was performed on lists of DMR-associated genes using WebGestalt [22].

**Assay for transposase-accessible chromatin (ATAC) sequencing**

For each sample, 50,000 cells were lysed, transposed, and PCR amplified as previously described [26], using the Illumina Nextera DNA kit. PCR products were purified using the Qiagen MinElute Kit (Qiagen), followed by a 1.8x AMPure XP bead selection (Beckman Coulter, Brea, CA, USA) to remove primer dimers. Libraries were sequenced on the Illumina NextSeq 500 using 2x75 bp paired-end runs and 1% PhiX spike-in. Reads were mapped to GRCh38 with Bowtie2 [23] and peaks were called using MACS2 [27]. Peak reads from each sample were normalized to the total number of aligned reads and principal component analysis was performed using Diffbind [28]. For peak annotation to nearby genes, differential peak analysis by Diffbind was rerun with MACS2 peaks in the 15q13.3 microdeletion region removed. To generate a list of differentially accessible genes, peaks were assigned first to promoters (2 kb region upstream of the transcriptional start site), then to gene bodies, and finally to the nearest gene (intergenic). For genes with multiple significant peaks, the peak with the most significant p-value was chosen for downstream gene-based analyses.

Homer motif enrichment analysis [29] was performed using a list of differentially open promoters and a custom motif file. The openness of each promoter, defined as the 2 kb region upstream of the transcriptional start site, was calculated from mapped reads using the surrounding 1 Mbp as background as previously described [30]. Reads in the 15q13.3 microdeletion region were doubled prior to calculating openness. A list of promoters with differential openness was generated using the following filters: 1) average openness in control samples > 2 and average openness in deletion samples < 2, or visa versa, and 2) absolute value difference in average openness in control samples - average openness in deletion samples >= 0.5. A custom list of transcription factors with known motifs was collected from existing databases and previous publications including Homer, Jaspar, ENCODE, and Taipale. Motif results were filtered for best match score >= 0.6 and p-value <= 10^-12^.
